# Human enterochromaffin cells and apical-out intestinal organoids as models for human sapovirus infection

**DOI:** 10.64898/2026.03.06.710023

**Authors:** Magdalena Neijd Segerstedt, Johan Nordgren, Berit Hammas, Jan Albert, Lennart Svensson, Marie Hagbom

## Abstract

Human sapovirus is increasingly recognized as a significant cause of acute viral gastroenteritis, but *in vitro* studies have been limited due to the previous lack of cell models. In this study, we optimized two novel *in vitro* models for sapovirus infection: the human enteroendocrine cell (EEC) line GOT1 and apical-out 3D human enteroids. Using 31 sapovirus-positive fecal samples, we compared these models with the EEC-derived HuTu80, and human enteroid monolayers, both previously used for sapovirus infection. Among the analyzed samples, genotypes GI.1 (36%), GII.1 (32%), and GII.3 (26%) were identified, with GI.1 exhibiting the highest fecal viral load. We demonstrated that GOT1 cells supported replication of more of the samples containing GI.1 sapovirus (9 of 11) compared to HuTu80 (8 of 11) and with a generally higher replication fold change. Moreover, GOT1 cells were able to support the replication of several GII.1 (3 of 10) and GII.3 (5 of 8) samples, unlike HuTu80 cells that only supported replication of one sample with GII.3 and none of the GII.1 sapoviruses. Given that organoids are considered a more physiologically relevant model than transformed cell lines, we established a sapovirus infection model using human apical-out 3D enteroids. Compared to previously used enteroid monolayers, the apical-out model supported replication of a higher number of samples with GI.1 and with higher replication fold change. In conclusion, these findings provide valuable insights for future *in vitro* studies of sapovirus infection and replication, which may contribute to a better understanding of sapovirus cell tropism and pathogenesis.

**Importance:** Understanding sapovirus biology requires efficient, reliable, and physiologically relevant *in vitro* models. By systematically comparing four infection models using clinical samples from three common sapovirus genotypes, this study contributes to important information about differences in cellular susceptibility between these *in vitro* models. We identified the GOT1 cell line as an efficient model for sapovirus replication and introduced apical–out enteroids as a sensitive organoid-based system for studying sapovirus GI.1. Together, the introduced models provide complementary platforms that can contribute to knowledge about sapovirus pathogenesis and cell tropism as well as provide guidance in selecting suitable *in vitro* models for future sapovirus research.

## Introduction

Members of the *Caliciviridae* family are the most common viral cause of acute gastroenteritis (AGE) worldwide, leading to numerous hospitalizations and deaths each year (1–5). While norovirus is the most well-known virus in this family, human sapovirus is also a significant cause of viral gastroenteritis, although usually with milder symptoms than norovirus (6). Sapovirus causes AGE across all age groups but is particularly prevalent among children, older adults, and immunocompromised individuals (7,8). Over the past five years, sapovirus has been identified as an important contributor to childhood diarrhea, especially following the introduction of the rotavirus vaccine (9–12). A study in the Nicaragua reported sapovirus as the second most detected viral pathogen in fecal samples from AGE patients, following norovirus (13). Furthermore, a meta-analysis of more than 100 studies across 35 countries found that sapovirus accounted for 3.4% of all AGE cases, emphasizing its global impact (14).

Sapoviruses are single-stranded, positive-sense, non-enveloped RNA viruses. Their genome consists of two open reading frames (ORFs) encoding non-structural proteins, a capsid protein, and several proteins with unknown functions. In some strains, a third ORF has been described, but its function remain unknown (15). Like norovirus, sapovirus exhibits high genetic diversity, with 19 genogroups (GI–GXIX) infecting a wide range of mammalian species (16,17). Among them, GI, GII, GIV, and GV infect humans, comprising 19 identified genotypes (GI.1–7, GII.1–8, GIV.1, and GV.1–2) (14,18,19).

Historically, the lack of reliable *in vitro* models has hindered research on sapovirus cell tropism and pathogenesis. Although sapovirus was first identified in 1976 (20), it was not successfully cultivated *in vitro* until year 2020, when Takagi *et al.* replicated and passaged the virus using the transformed duodenal EEC cell line HuTu80 (21). Therefore, much remains unknown about sapovirus cell tropism, pathogenesis, and host immunity. Following its successful replication in HuTu80 cells, sapovirus was subsequently shown to infect monolayers of human intestinal organoids from all three segments of the small intestine but not monolayers derived from colonoids (22). A key advantage of human intestinal organoids is that they are non-transformed and differentiate into multiple epithelial cell types, which represent the physiologically relevant target cells for enteric virus infections (23), making them more accurate, representative and translational, compared to transformed cell lines. Although enteroid monolayers are commonly used in infection studies (24), their effectiveness depends on achieving a fully confluent and stable monolayer throughout the experiment. As an alternative, Co *et al.* recently proposed using three-dimensional (3D) organoids in suspension (apical-out) for experimental infections (25), a model proven efficient for norovirus (26). In this study, we established the apical-out 3D enteroid model for sapovirus infection studies and compared infection and replication efficacy to monolayers. Also, since HuTu80 cells originate from EECs, a duodenal adenocarcinoma of L-cell type (ATCC HTB-40), we investigated whether another EEC cell line, GOT1, could serve as a potential model for sapovirus infection. GOT1 cells, established in year 2001 from a human neuroendocrine tumor metastasized to the liver (27), are serotonin-producing EC cells, a subset of EECs originally found in the intestine (28). Previous studies have shown that rotavirus and its enterotoxin NSP4 can infect respectively stimulate GOT1 cells to trigger serotonin secretion (29). Similarly, human norovirus has been demonstrated to infect EECs in the small intestine (30), suggesting that sapovirus may also exhibit enteroendocrine cell tropism.

Here, we compare replication of three human sapovirus genotypes across the four *in vitro* models; HuTu80, GOT1, enteroid monolayers and apical-out 3D enteroids. Our findings reveal distinct differences in susceptibility and replication efficiency between the models. Notably, the GOT1 cell line supported replication of highest number of sapovirus-positive fecal samples and with higher replication, indicating its potential as a suitable and efficient model for sapovirus research. Furthermore, we demonstrated that apical-out 3D enteroids were more susceptible to sapovirus GI.1 infection than monolayers, highlighting their potential for future sapovirus infection studies.

## Results

### Higher viral load in fecal samples with sapovirus genotype GI.1

A total of 31 sapovirus-positive fecal samples were collected from individuals seeking healthcare for AGE during 2021 and 2022. Among these, 11 samples (36%) were identified as genotype GI.1, 10 (32%) as GII.1, and 8 (26%) as GII.3 (Figure 1A). Two samples (6%) were un-typeable. Quantification by qPCR showed that the viral load of samples containing sapovirus GI.1 had highest median fecal viral load (3.3 × 10¹⁰ genome copies per gram of feces), compared to 2.7 × 10⁹ for GII.1 (p = 0.0472) and 6.1 × 10⁸ for GII.3 (p = 0.0002). The un-typeable samples had a median viral load of 4.5 × 10⁸ genome copies per gram (Figure 1B).

**Figure 1.**
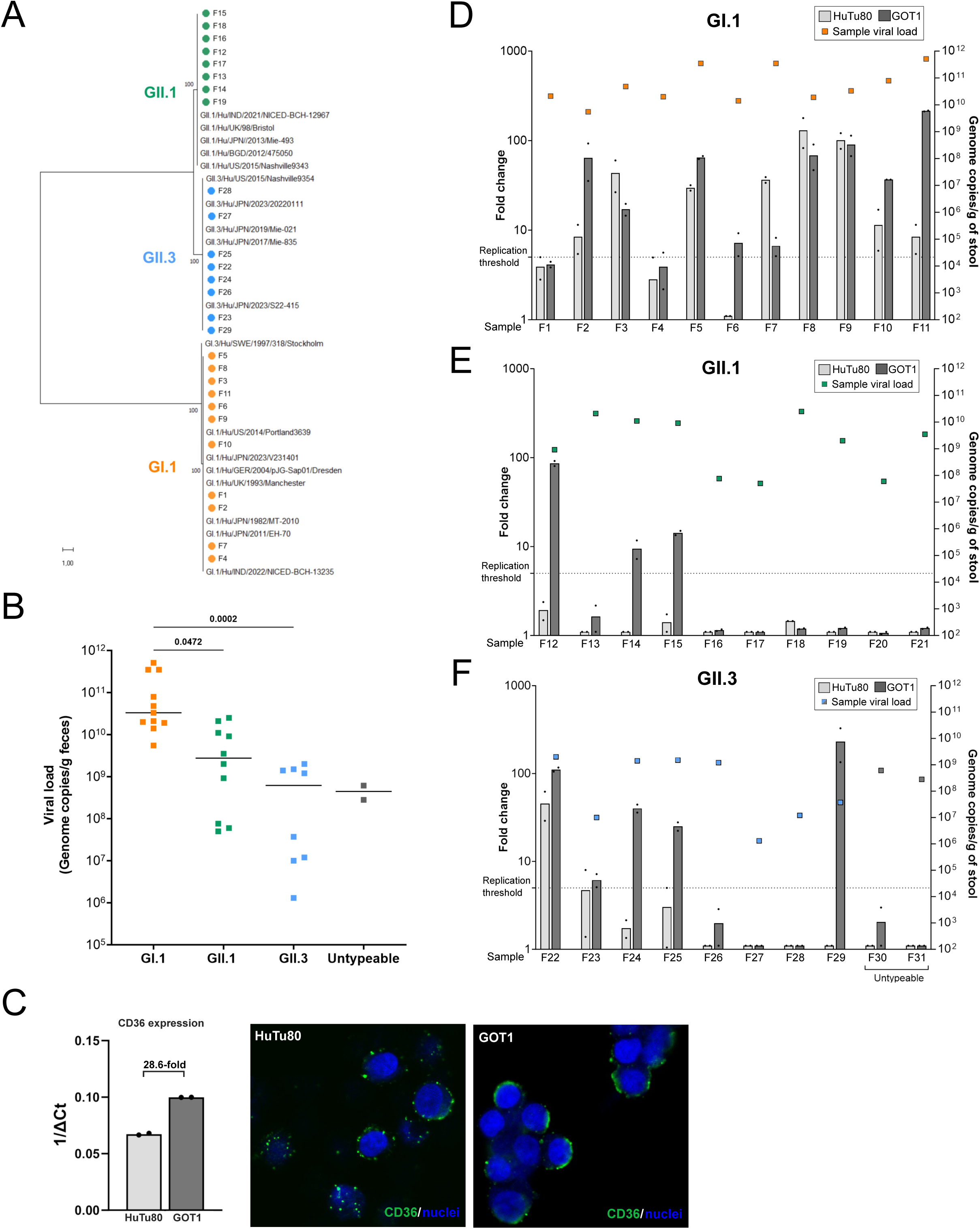
A) Phylogenetic tree of sapovirus samples (n = 27), color coded by genotype, and reference strains. The tree was constructed with Maximun likelihood algorithm using Kimura 2-parameter model (gamma distributed with invariant sites) and 1000 bootstrap replicates. Scale bar for branch lengths shows number of substitutions per site. Samples F20, F21, F30 and F31 were excluded in the analysis due to incomplete sequences. Reference strains: GII.1/Hu/IND/2021/NICED-BCH-12967 (GenBank: LC851686), GII.1/Hu/JPN/2013/Mie-493 (GenBank: LC816490), GII.1/Hu/BGD/2012/475050 (GenBank: MW088928), GII.1/Hu/UK/1998/Bristol (GenBank: AJ249939), GII.3/Hu/JPN/2023/20220111 (GenBank: LC848995), GII.3/Hu/JPN/2023/S22-415 (GenBank: LC849000), GII.3/Hu/JPN/2019/Mie-021 (GenBank: LC816544), GII.3/Hu/JPN/2017/Mie-835 (GenBank: LC816541), GI.1/Hu/US/2014/ Portland3639 (Genbank: MG012399), GI.1/Hu/IND/2022/NICED-BCH-13235 (GenBank: LC851695), GI.1/Hu/JPN/2011/EH-70 (GenBank: LC504316), GI.1/Hu/JPN/2023/V231401 (GenBank: LC848977), GI.1/Hu/JPN/1982/MT-2010 (GenBank: HM002617), GII.1/Hu/US/2015/Nashville9343 (GenBank: MG012444), GI.1/Hu/GER/pJG-Sap01/Dresden (GenBank: AY694184), GI.3/Hu/SWE/1997/318/Stockholm (GenBank: AF194182), GII.3/Hu/US/2015/Nashville9354 (GenBank: MG012419), GI.1/Hu/UK/1993/Manchester (GenBank: X86560). Abbreviations: Hu: Human, IND: India, JPN: Japan, US: United States, BGD: Bangladesh, GER: Germany, SWE: Sweden, UK: United Kingdom. **B)** Viral load (genome copies/gram of feces) and genotype of 31 sapovirus positive samples. Each colored square corresponds to one sample, and the median viral load is shown for each genotype. p-value, determined by Kruskal-Wallis test, is displayed. **C)** Detection of CD36 by gene expresison analysis and immunofluorescence staining in GOT1 and HuTu80. Gene expression of CD36 is displayed as inverted ΔCt value (1/ ΔCt), with ΔCt calculated by subtracting the Ct-value for CD36 to the Ct of the house-keeping gene GAPDH. The fold difference in expression between HuTu80 and GOT1 is shown above the bars. **D - F)** Replication of sapovirus samples in HuTu80 and GOT1. Left y-axis represents the fold change (FC) of virus at 72 hours post infection (hpi) compared to 2 hpi. The two biological duplicates per infection experiment are shown as black dots. A mean FC of at least 5-fold (dotted line) was considered successful replication and samples with a mean FC of less than 1 were set to 1.1 FC for visualization purposes in the graph. The sample viral load, expressed as genome copies per gram of feces, is shown as colored squares. GAPDH; Glyceraldehyd-3-fosfatdehydrogenas

### Sapovirus replicates efficiently in GOT1 enterochromaffin cells

The enteroendocrine HuTu80 cell line has previously been shown to support sapovirus replication (21). To further evaluate different *in vitro* models for sapovirus, we assessed the susceptibility of the EC cell line GOT1 and compared it to HuTu80. First, the presence of CD36 was assessed by gene expression analysis and immunofluorescence staining in both HuTu80 and GOT1 since it has been identified as an essential cell surface protein for human sapovirus infection (31). Expression of CD36 was found in both HuTu80 and GOT1, with 28.6-fold higher expression in GOT1 cells, and was detected on the cell surface of both cell lines by immunofluorescence staining (Figure 1C).

Next, all sapovirus samples (F1 – F31) were tested in both GOT1 and HuTu80 (Figure 1D-F). The fold change (FC) in viral copies at 72 hours post-infection (hpi) compared to 2 hpi was determined by qPCR. Replication in HuTu80 (FC ≥ 5 at 72 hpi) was observed in 8 of 11 samples containing GI.1 and 1 of 11 GII.3 samples, whereas no replication was observed for the samples containing GII.1, even at high viral loads (>10⁹ viral copies per gram of feces). The FC at 72hpi for GI.1 sapoviruses were between 8.4 and 130.6 (median 33.1-fold), while the sample with GII.3 showed a FC of 45.6-fold (Figure 1D-F). All eight samples with sapovirus GI.1 that had replicated in HuTu80 also replicated in GOT1 cells, with FC at 72 hpi ranging from 6.8-fold to 214.5-fold (median 64.5-fold). GOT1 also supported replication of the sample with sapovirus GII.3 that had replicated in HuTu80 (F22) with a FC of 111.2-fold. Moreover, GOT1 cells supported replication of sapovirus samples that did not replicate in HuTu80 cells, including one GI.1 sapovirus (36.8-fold), four samples with GII.3 sapovirus, ranging from 6.1-fold to 231.1-fold, and three samples with GII.1, ranging from 9.5-fold to 86.1-fold. In summary, 9 of 11 samples with GI.1 sapovirus, 3 of 10 samples with GII.1, and 5 of 8 GII.3 samples replicated in GOT1 cells (Figure 1D-F). None of the un-typeable viruses replicated in either HuTu80 or in GOT1 (Figure 1F).

Samples with replicating sapovirus had higher fecal viral load than non-replicating virus samples in both HuTu80 (p = 0.0002) and in GOT1 (p = 0.0495) (Figure 2A), but no correlation was observed between fecal viral load and replication FC in either HuTu80 (p = 0.22) or GOT1 (p = 0.98) (Figure 2B). To further evaluate the novel GOT1 model, infection experiments were repeated in three independent experiments for one virus sample of each genotype. Replication was observed in all three experiments and for all three virus samples, with variation in mean FC ranging between 44 – 112 (GI.1), 34 – 86 (GII.1) and 111 – 744 (GII.3) (Figure 2C). Moreover, replication kinetics in GOT cells demonstrated that viral replication (FC ≥ 5) of GI.1 and GII.3 occurred already at 12 hpi. By 1 day post–infection (dpi), all three genotypes (GI.1, GII.1 and GII.3) exhibited a marked increase in FC. For GI.1 and GII.1, replication FC thereafter plateaued after 1 dpi and remained relatively constant throughout the experiment, until terminated at 7 dpi. In contrast, replication FC for GII.3 continued to rise steadily throughout the 7–day period (Figure 2C).

**Figure 2.**
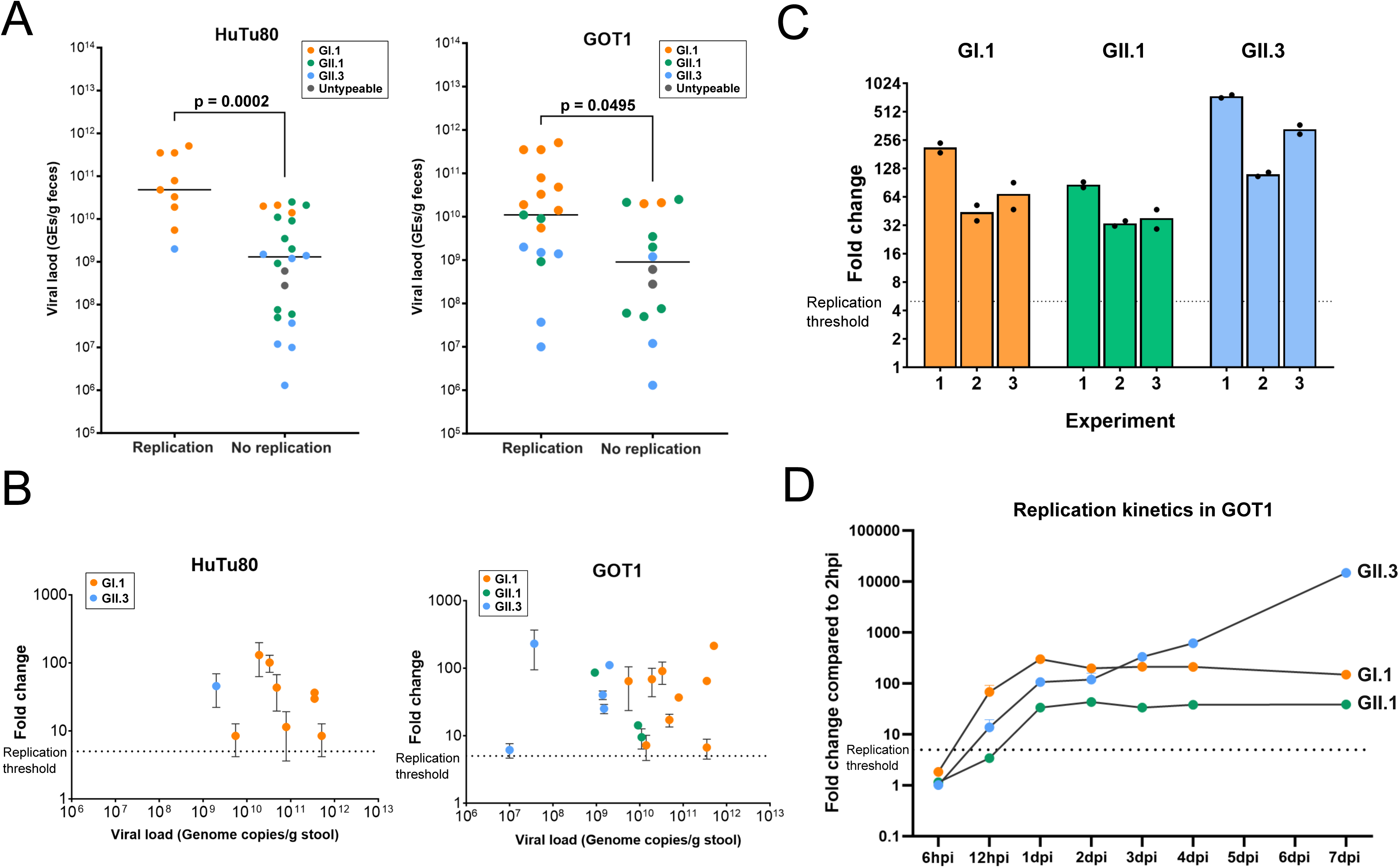
A) Viral load for replicating and non-replicating samples in HuTu80 (left) and GOT1 (right), color coded by genotype. p-value, calculated by Mann-Whitney U test, is displayed. **B)** Correlation analysis of replication FC and sample viral load in HuTu80 (left) and GOT1 (right), determined by Spearman r. Left y-axis represents the fold change (FC) of virus at 72 hours post infection (hpi) compared to 2hpi. The sample viral load, displayed as genome copies per gram of stool, is shown on x-axis. Error bars show standard deviation in FC between the two biological replicates of the infection experiment. **C)** Replication of sapovirus GI.1, GII.1 and GII.3 in GOT1 cells in three independent experiments. y-axis shows the fold change (FC) of virus at 72 hours post infection (hpi) compared to 2hpi. The two biological duplicates per infection experiment are shown as black dots. A mean FC of at least 5-fold (dotted line) was considered successful replication. **D)** Replication kinetics of GI.1, GII.1 and GII.3 in GOT1 cells, presented as a fold change in viral RNA at different time points compared to at 2 hpi. Dotted line represents replication threshold of FC=5.

### Apical-out 3D enteroids as an alternative model for sapovirus infection

As previously reported, sapovirus GI.1 can infect enteroid 2D monolayers (22). Since enteroids provide a more physiologically relevant model than transformed cell lines, this model is more suitable for pathogenesis studies. As a potential sapovirus infection model, we generated apical-out 3D enteroids in suspension as previously described by Co et al. (25). The reversion of epithelial polarity from basal-out to apical-out conformation was visualized by fluorescently labeled lectin detecting apical glycoproteins. In contrast to enteroids in Matrigel, all enteroids in suspension successfully had reversed the polarity on day 7 after removing extracellular matrix proteins (Figure 3A). CD36 was detected in both differentiated monolayers and apical-out enteroids, with 30.6-fold more mRNA expression in monolayers (Figure 3B). In addition, the expression of differentiation-associated genes was assessed relative to undifferentiated enteroids in both monolayers and apical-out enteroids. Differentiation led to upregulation of the enterocyte marker sucrase isomaltase (SI) and the goblet cell marker mucin-2 (MUC2) in both models. In contrast, the stem cell marker Leucine-rich repeat-containing G-protein coupled receptor 5 (LGR5) was downregulated in differentiated monolayers but showed a slight increase in apical-out enteroids. Expression of the enteroendocrine cell marker chromogranin A (CHGA) was upregulated in apical-out enteroids, whereas its expression in differentiated monolayers varied between experiments (Figure 3C).

**Figure 3.**
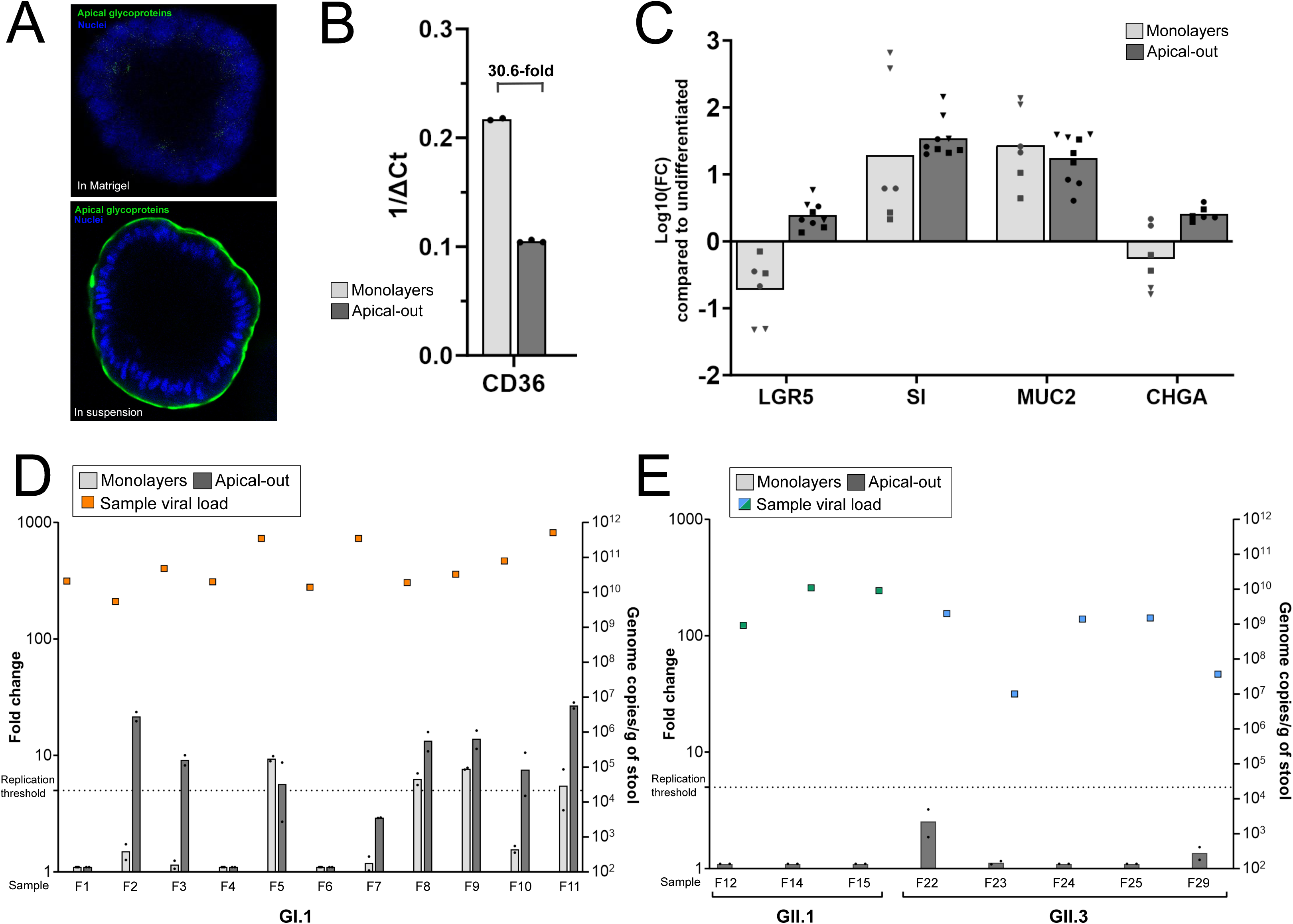
A) FITC-lectin staining of apical glycoproteins in differentiated enteroids in Matrigel (basal-out) and in suspension (apical-out). **B)** Gene expression of CD36 in monolayers and apical-out enteroids, displayed as inverted ΔCt value (1/ ΔCt), with ΔCt calculated by subtracting the Ct-value for CD36 to the Ct of the house-keeping gene GAPDH. The fold difference in expression between monolayers and apical-out enteroids is shown above the bars. **C)** Gene expression analysis of the differentiation-associated genes LGR5, SI, MUC2 and CHGA in enteroid monolayers and apical-out enteroids in three independent experiments with 2-3 biological replicates each, shown as different symbol shapes. Data is shown as log10 (fold change) in expression at 5 days of differentiation compared to undifferentiated enteroids, calculated by the 2^-ΔΔCt^ (Delta-Delta Ct) method using GAPDH as housekeeping gene. In one of the experiments in apical-out enteroids, no CHGA expression was detected and was therefore not included in the graph. **D - E)** Replication of sapovirus GI.1 samples in enteroid monolayers and apical-out enteroids (D) and GII.1 and GII.3 in apical-out enteroids (E). Left y-axis represents the fold change (FC) of virus at 72 hours post infection (hpi) compared to 2 hpi. The two biological duplicates per infection experiment are shown as black dots. A mean FC of at least 5-fold (dotted line) was considered successful replication and samples with a mean FC of less than 1 were set to 1.1 FC for visualization purposes in the graph. The sample viral load, displayed as genome copies per gram of stool, is shown as colored squares. GAPDH; Glyceraldehyd-3-fosfatdehydrogenas, LGR5; Leucine-rich repeat-containing G-protein coupled receptor 5, SI; sucrase isomaltase, MUC2; mucin-2.

Further on, we evaluated sapovirus replication in both enteroid monolayers and apical-out enteroids by testing all 11 samples with GI.1 that were previously tested in the transformed cell lines (Figure 3D). Four of the 11 virus samples replicated in monolayers, with a FC ranging from 6.3 to 9.4 (median 7.2-fold). In the apical-out model, 7 of 11 samples replicated, exhibiting FC ranging from 5.7 to 26.9 (median 13.4-fold). Moreover, since the apical-out model supported replication of more virus samples than monolayers, we proceeded with investigating whether this 3D model could support replication of other genotypes than GI.1 by testing the samples containing GII sapovirus that had replicated in the transformed cell lines, including three GII.1 (F12, F14 and F15) and five GII.3 samples (F22 – F25 and F29). None of these samples replicated in the apical-out model (Figure 3E).

## Discussion

Efficient, reliable and representative *in vitro* models are essential for understanding cell tropism, infectiousness, and pathogenesis of viruses. In this study, we evaluated four *in vitro* models for human sapovirus replication, using 31 fecal samples of three different sapovirus genotypes. The genotypes were distributed between GI.1 (36%), GII.1 (32%), and GII.3 (26%), which are all common globally (32,33). Here, fecal samples with GI.1 exhibited highest fecal viral load, as quantified by qPCR, which may indicate a greater replication efficiency for this genotype *in vivo*. Supporting this, a birth cohort in Nicaragua showed that sapovirus GI.1 was more common in symptomatic infections compared to asymptomatic, suggesting that this genotype may be more pathogenic than others (34).

Infectivity of all 31 sapovirus-positive fecal samples were tested in the transformed cell lines HuTu80 and GOT1, where we observed differences in susceptibility and replication between the models. For GI.1, both models supported replication, though GOT1 cells supported replication of more samples than HuTu80 (82% vs. 70%). In contrast, HuTu80 cells only supported replication of one GII.3 sample, whereas GOT1 cells supported replication of more than half of the samples (63%). Notably, GOT1 was the only model that supported replication of GII.1, with 30% of samples replicating. These results clearly demonstrate that the GOT1 cell line is a functional cell model for sapovirus replication and in our hands more sensitive to sapovirus infection compared to HuTu80 cells. The failure of GII.1 to replicate in HuTu80 cells contrasts findings of Takagi *et al.* (35), who reported successful replication of all three genotypes in HuTu80 cells, including a GII.1 sample with a lower viral load than those tested in our study. Takagi *et al.* have demonstrated that high passage number of HuTu80 cells, preferably more than 140 passages, and the use of the bile acid Taurocholate (TauCA) during infection experiments enhances the replication efficacy in the model (35), which may explain the differences to our replication data. However, in our hands, using TauCA instead of GCDCA did not enhance replication in either HuTu80 or GOT1 cells (data now shown). Interestingly, a recent publication also highlighted failed replication of sapovirus in HuTu80 cells, and demonstrated that the cell line Caco-2MC, a Cas9-expressing Caco-2 cell line (Caco2/Cas9) (36), was highly susceptible to sapovirus, arguing that it is a robust alternative to the HuTu80 cell line (37).

In this study, we confirmed that enteroid monolayers are susceptible to sapovirus GI.1, as reported by Euller-Nicolas *et al.* in 2023 (22). Furthermore, when comparing enteroid models, we demonstrated that apical-out 3D supported replication of more samples with GI.1 than enteroid monolayers. 7 of 11 GI.1 samples (64%) replicated in the apical-out model, and only 4 of 11 (36%) replicated in monolayers, and replication efficiency, measured by FC, generally being higher in the apical-out model. Several biological and structural differences may account for this enhanced permissiveness, including the increased accessible cell surface area for virus attachment in the 3D organoids and the ability of apical-out enteroids to sustain stemness after differentiation, as shown in this study. We further tested whether sapovirus from the GII genogroup could infect the apical-out enteroids. Three GII.1 and five GII.3 sapovirus samples that had shown replication in GOT1 cells were tested in the model but failed to replicate. This suggests a genotype dependent susceptibility in the enteroid model and further studies, including additional genogroups and genotypes, are needed to better understand the genotype-specific replication patterns in enteroids.

CD36 has recently been proposed as an essential host factor for sapovirus infection (31), and our findings provide additional support for this since CD36 expression was detected in all four model systems. Notably, GOT1 cells, which supported sapovirus replication of the highest number of samples and generally exhibited the highest FC, displayed markedly higher CD36 expression compared to HuTu80 cells, suggesting that CD36 expression may contribute to cellular permissiveness. However, this association was not consistent when comparing the enteroid models. Although CD36 expression was substantially higher in enteroid monolayers than in apical-out enteroids, monolayers supported replication in fewer samples and with lower FC. This discrepancy indicates that while CD36 appears to be necessary for sapovirus infection, its expression level alone is insufficient to explain differences in replication efficiency.

In summary, our results show that the GOT1 cell line was the most efficient infection model for human sapovirus, supporting replication of the highest number of sapovirus samples, with generally the highest FC in viral replication, across all four infection models and the three tested sapovirus genotypes. The apical-out 3D enteroid model was not susceptible to the GII genogroup but was able to support replication of the majority of GI.1 samples, in contrast to enteroid monolayers, making it a valuable model for sapovirus research as a more sensitive physiologically relevant model system. The successful replication of sapovirus in both HuTu80 and GOT1 cells suggests a potential sapovirus cell tropism for EEC cells. By using the enteroid model, future studies could focus on localizing sapovirus within specific cell types to further characterize the cell tropism. Our findings are of value for future *in vitro* studies of sapovirus infection and may contribute to a deeper understanding of sapovirus cell tropism and pathogenesis.

## Material and methods

### Fecal sample preparation

Sapovirus positive fecal samples (Ct < 29) were obtained from the clinic. Each sample was diluted as a 10 % suspension in organoid basal media (Gibco™ Advanced Dulbecco’s Modified Eagle Medium/F12 supplemented (DMEM/F12) with 10 mM HEPES, 1X GlutaMAX, 20 µg/ml Gentamicin). After centrifugation for 5 minutes at 13 000 x g, the suspension was aliquoted and kept at −80°C until further use. 140 μl of the 10% fecal suspension was lyzed in 560 μl AVL (Qiagen) for genotyping and viral quantification.

### Transformed cell line cultures

The duodenal adenocarcinoma cell line HuTu80 (< 50 passages) was cultured in Gibco™ Dulbecco’s Modified Eagle Medium (DMEM) containing 20 µg/ml Gentamicin (Gibco) and 10% fetal calf serum (FCS), at 37°C and 5% CO_2_. The cells were split every third day by trypsinization in 0.25% Trypsin-EDTA (Gibco). For experiments, the cells were plated in 48 or 96 well plates.

The enteroendocrine cell line GOT1 (< 115 passages) was cultured in Gibco™ RPMI media containing 1X Minimum Essential Medium (MEM) non-essential amino acids (MEM NEAA) (Gibco), 20 µg/ml Gentamicin, 1 mM Sodium pyruvate (Gibco) and 2.4X GlutaMAX™ (Gibco) and 10% FCS, at 37°C and 5% CO_2_. The cells were split every 10-14 days by trypsinization in 0.25% Trypsin-EDTA. For experiments, the cells were plated in Nunc™ Nunclon 48 well plates (ThermoFisher).

### Growth and maintenance of enteroids

Enteroids were grown in 3D Matrigel® Matrix domes (Corning®) and human STEMCELLTechnologies™ IntestiCult™ Organoid Growth Media supplemented with 1:1 ratio of Organoid Supplement media (STEMCELL Technologies, UK) and 20 µg/ml Gentamicin (complete OGM) at 37°C and 5% CO_2_. Media was changed every second to third day and the cells were passaged once a week in a 1:2 ratio. For passaging, the Matrigel dome was dissolved in STEMCELL Technologies™ Gentle Cell Dissociation Reagent (GCDR), and the collected organoids were washed in organoid basal media containing 10% FCS. After centrifugation, the pellet of organoids was resuspended in Matrigel and added dropwise (50 µl) to form new Matrigel domes. The domes were let solidify before complete OGM was added to each well.

### Generation of enteroid monolayers

To obtain and differentiate enteroids as 2D monolayers, Matrigel domes were dissolved in Gibco™ Versene Solution, washed in organoid basal media and the obtained cells resuspended in 0.05% Trypsin-EDTA (Gibco). Trypsin was inactivated in organoid basal media with 10% FCS, and the cell solution was vigorously pipetted until single cells were obtained. After filtering through a 40 µm filter, and followed by centrifugation, the single cells were resuspended in complete OGM containing 10 µM ROCK inhibitor and transferred to each well of a 96 well plate, pre-coated with CellAdhere™ Type I Collagen Bovine Solution (STEMCELL Technologies, UK). After 2 days of culture in complete OGM at 37°C and 5% CO_2_, the growth media was changed to STEMCELL Technologies™ IntestiCult™ Organoid Differentiation Media supplemented with 1:1 ratio of Organoid Supplement media (STEMCELL Technologies, UK) and 20 µg/ml gentamicin (complete ODM) to initiate differentiation for 4 days before infection.

### Infection of transformed cell lines and enteroid monolayers

The 10 % fecal samples were diluted to a 1 % suspension in serum free DMEM (HuTu80), organoid basal media (enteroids) or RPMI supplemented 1X Minimum Essential Medium (MEM) non-essential amino acids (MEM NEAA), 20 µg/ml Gentamicin, 1 mM Sodium pyruvate and 2.4X GlutaMAX™ (GOT1) containing 500 μM of Sigma-Aldrich glycochenodeoxycholic acid (GCDCA). The cells were washed once in appropriate serum free media before the 1 % fecal suspension was added to the cells followed by incubation at 37°C for 2 hours. The cells were washed twice in serum free media followed by addition of media containing FCS and 500 μM of GCDCA. The 2-hour samples were lyzed in AVL buffer (Qiagen) according to manufacturer’s protocol, and remaining wells were incubated until 72 hpi before lyzed for quantification of viral RNA by qPCR. For infection kinetics in GOT1 cells, samples were also lyzed at 6 hpi, 12 hpi, 1 dpi, 2 dpi, 3 dpi, 4 dpi and 7 dpi.

### Generation and infection of apical-out enteroids

To obtain apical-out enteroids, the Matrigel was dissolved in Versene solution and organoids collected in tubes. The organoids were washed twice in organoid basal media followed by centrifugation at 200 x g at 4 °C. At the last washing step, the cell pellet was resuspended into complete OGM and transferred to a 48 well plate and incubated at 37 °C and 5% of CO_2_. After 48 hours, the free-floating apical-out enteroids were collected in Eppendorf tubes and centrifugated at 90 x g for 3 minutes. The supernatant was discarded, and organoids resuspended in complete ODM to initiate differentiation.

On day 4 of differentiation, the organoids were collected in Eppendorf tubes and washed once in organoid basal media and thereafter resuspended in 1% fecal suspension diluted in organoid basal media containing 500 μM of GCDCA. The enteroids were incubated in one well for 2 hours at 37°C and then collected in Eppendorf tube and washed three times in organoid basal media. In the last step, enteroids were resuspended in complete ODM containing 500 µM GCDCA and evenly distributed into wells. The 2-hour wells were lyzed in AVL buffer and remaining wells incubated for 72 hours before lyzed for quantification of viral RNA by qPCR.

### RNA extraction and cDNA synthesis

Viral RNA was extracted using QIAamp® Viral RNA Mini Kit (QIAGEN) spin column and cell RNA for gene expression analysis was extracted using RNeasy mini kit (QIAGEN) according to the manufacturer’s protocol. RNA was transcribed to cDNA using High-capacity cDNA kit (Applied Biosystems) and stored in −80°C until PCR analysis.

### Determination of sapovirus replication by qPCR

To measure viral replication, the FC of the amount of viral RNA in samples at 72hpi was compared to 2hpi. Amount of viral RNA was determined by qPCR using iTaq™ Universal Probes Supermix, 200 nM probe (SaV124TP), 400 nM of two forward primers (SaV124F and SaV1F) and 400 nM reverse primer (SaV1245R) (38). qPCR run conditions started with a denaturation step at 95°C for 5 minutes followed by 15 seconds at 95°C and 60°C for 1 minute for 44 cycles.

### Gene expression analysis by qPCR

Expression of CD36 and genes associated with stemness and differentiation in enteroids was determined by qPCR. The qPCR analysis was performed in technical duplicates on a CFX96™ Real-Time System (Bio-Rad, USA) using SsoAdvanced Universal SYBR Green Supermix (Bio-Rad Laboratories, USA) and SYBR probes (Supplementary table 1) according to the manufacturer’s instruction. Expression of the target genes were normalized to the level of the housekeeping gene Glyceraldehyd-3-fosfatdehydrogenas (GAPDH) (target Ct – GAPDH Ct). For determining differentiation in enteroids, the 2^-ΔΔCt^ method was used to calculate the FC in expression in differentiated compared to undifferentiated samples.

### Genotyping and quantification of sapovirus in fecal samples

The cDNA generated from the 10% fecal suspension was used for viral quantification and genotyping. Quantification of virus by qPCR was performed as described above, where plasmids with synthesized sapovirus sequence of different dilutions were used to generate a standard curve. Genotyping was performed using a PCR amplification strategy previously described by Moraes *et. al* (39), using iTaq^TM^ DNA Polymerase PCR kit. The successfully amplified PCR products were sent to Macrogen Europe for sanger sequencing, and the obtained data of the forward and reverse strands were manually analyzed and aligned in BioEdit 7.2.5 to create a consensus sequence. A phylogenetic tree was constructed based on the consensus sequences and generated in MEGA11 (Molecular Evolutionary Genetics Analysis) software version 11.0.8 using maximum likelihood algorithm with Kimura 2-parameter model (gamma distributed with invariant sites) and 1000 bootstrap replicates.

### Immunofluorescence staining

The cells were fixed in 4% formaldehyde for 15-30 minutes in RT or in 4°C over-night followed by two washing steps with phosphate buffer saline (PBS). Apical-out enteroids were stained in low-binding Eppendorf tubes. Between each step of the staining protocol, the 3D enteroids were centrifugated for 3 minutes at 90 x g followed by aspiration of supernatant. Blocking was performed with 3% bovine serum albumin (BSA) (Sigma-Aldrich, #A2153) in PBS for 60-90 minutes at RT. After two washes, primary antibody 1:200 diluted in 1% BSA in PBS was added and incubated overnight at 4°C for enteroids or 2 hours at RT for transformed cell lines. Samples were washed five times in PBS before adding secondary antibody 1:200 diluted in 1% BSA in PBS and incubated 4 hours (enteroids) or 2 hours (GOT1 and HuTu80) at RT followed by five washed in PBS. DAPI 1:1000 diluted in PBS was added and incubated 5-10 minutes at RT followed by three washes. After the last washing step, apical-out enteroids were resuspended in PBS and added to Nunc™ Lab-Tek™ Chamber Slide System (Thermo Scientific, #155411PK). The antibodies are summarized in supplementary table 1. Immunofluorescence staining’s were visualized in Zeiss LSM700 upright confocal microscope or LSM700 inverted confocal microscope. Color balance in the images was adjusted in Fiji ImageJ version 1.53c.

### Statistical analyses

Statistical analyses were performed using GraphPad Prism version 10.0.2 (232). Due to the small samples size, Shapiro-Wilk statistical was used to determine normality, showing that none of the results were normally distributed. Hence, all statistical analyses were performed using non-parametric tests, all of which are indicated in the figure legends.

## ACKNOWLEDGMENTS

This study was supported by the Swedish Research Council (2023-02720). Our sincere gratitude to Claudia Beck Eichler Jonsson at Karolinska Institute for organizing logistics behind sample storage and transportation to Linköping University.

